## Supplemental Information for "Subjective value and decision entropy are jointly encoded by aligned gradients across the human brain"

### Overview from NARPS

The NARPS team collected collected fMRI data on two versions of a mixed gambles task [Tom et al., 2007]. On each trial, a mixed gamble was presented (one gain amount, one loss amount) and the participants decided whether to accept the gamble or not. Each participant was assigned to one of two conditions: In the equal indifference condition, the matrix of gambles included potential gains twice the range of potential losses [Tom et al., 2007]; in the equal range condition, the matrix included an equal range of potential gains and losses [De Martino et al., 2010]. For completeness, below we present the original experimental protocol, data description, MRI scanning protocols, fmriprep preprocessing output, and the original ex-ante hypotheses from the NARPS team. Alternatively, this information is also available on the NARPS website http://www.narps.info, the data descriptor [Botvinik-Nezer et al., 2019], and/or the ensuing publication [Botvinik-Nezer et al., 2020].

### Experimental protocol and instructions (NARPS)

Upon arrival, as soon as they signed the consent forms, participants were endowed with 20 ILS ( $5.5 USD) cash payment for their participation. The experimenter explained that the money is theirs to keep, and is part of the full amount they would receive at the end of the experiment. Next, the participants received general instructions regarding behavior inside the scanner and performed a shortened version of the full task (i.e. a demo).

Then, participants entered the MRI scanner. Participants completed four runs of the mixed gambles task, each consisting of 64 trials. On each trial, a mixed gamble was presented, entailing a 50/50 chance of gaining one amount of money or losing another amount. Possible gains ranged from 10-40 ILS (in increments of 2 ILS) or 5-20 ILS (equal indifference and equal range conditions, respectively) and possible losses ranged from 5-20 ILS (in increments of 1 ILS). All 256 possible combinations of gains and losses were presented across the four runs.

Stimulus presentation was similar to Tom et al. [2007]. Timing of all stimuli and response events were computed using Matlab 2014b and the Psychtoolbox [Pelli, 1997, Brainard and Vision, 1997] on an Apple MacBookPro running Mac OS X Yosemite version 10.10.5 (Apple Computers, Cupertino, CA). The timing and order of stimulus presentation was optimized for estimation efficiency using a tailored code from the creators of neuropowertools [Durnez et al., 2016]. Due to the fact that this experiment involved only one trial type, which is beyond the scope of the existing tool, a custom solution was required. The efficiency calculations assumed a 32s HRF, a 1s TR and a truncated exponential distribution of ITIs (min=6s, max=10s mean=7s, lambda was extrapolated from these parameters), the minimal ITI encompass a potential trial duration of 4s and 2s intermission.

To even the gambles between different runs, the full matrix of gambles was divided into 16 4X4 sub matrices, which were independently scrambled and allocated to the different runs. This procedure facilitated the overall similarity between runs. Eight different onsets were created using this procedure for each experimental condition.

As in Tom et al. [2007], participants were asked to evaluate whether or not they would like to play each of the gambles presented to them (strongly accept, weakly accept, weakly reject or strongly reject). They were told that one trial from each of the runs would be selected at random, and if they had accepted that gamble during the task, the outcome would be decided with a coin toss; if they had rejected the gamble, then the gamble would not be played.

Following imaging, participants were presented with questionnaires regarding gambling attitudes and made a number of choices involving hypothetical gambles.

### Data from NARPS

119 healthy participants completed the experiment ($n = 60$ from the equal indifference group and $n = 59$ from the equal range group). Nine participants were excluded prior to fMRI analysis based on pre-registered exclusion criteria: Five did not show a significant effect of both gains and loses on their choices (Bayesian logistic regression, $p < 0.05$; reflecting a lack of understanding of the task) and four missed over 10% of trials (in one or more runs). Data of two additional participants is currently under QA. Thus, at least 108 participants will be included in the final dataset sent to the analysis teams ($n = 54$ from the equal indifference group and $n = 54$ from the equal range group).

### MRI scanning protocols (NARPS)

MRI was performed on a 3T Siemens Prisma scanner at Tel Aviv University. MRI scanning included the following acquisitions:

**Structural MRI** High-resolution T1w structural images were acquired using a magnetization prepared rapid gradient echo (MPRAGE) pulse sequence with the following parameters: TR = 2530 ms, TE = 2.99 ms, FA = 7, FOV = 224 $\times$ 224 mm, resolution = 1 $\times$ 1 $\times$ 1 mm.

**Functional MRI** Whole-brain FMRI data were acquired using echo-planar imaging with multi-band acceleration factor of 4 and parallel imaging factor (iPAT) of 2, TR = 1000 ms, TE = 30 ms, flip angle = 68 degrees, in plane resolution of 2X2 mm 30 degrees of the anterior commissure-posterior commissure line to reduce the frontal signal dropout, with slice thickness of 2 mm and a gap of 0.4 mm between slices to cover the entire brain.

Data were converted to NIFTI format using dcm2nii [RRID:SCR_014099, Li et al., 2016] and transformed into the Brain Imaging Data Structure (BIDS) [Gorgolewski et al., 2016].

### fMRI preprocessing (NARPS)

Results included in this manuscript come from preprocessing performed using *fMRIPprep* 1.1.6 (Esteban et al. [2018b]; Esteban et al. [2018a];RRID:SCR_016216), which is based on *Nipype* 1.1.2 (Gorgolewski et al. [2011]; Gorgolewski et al. [2018]; RRID:SCR_002502).

**Anatomical data preprocessing** The T1-weighted (T1w) image was corrected for intensity non-uniformity (INU) using `N4BiasFieldCorrection` [Tustison et al., 2010, ANTs 2.2.0], and used as T1w-reference throughout the workflow. The T1w-reference was then skull-stripped using `antsBrainExtraction.sh` (ANTs 2.2.0), using OASIS as target template. Brain surfaces were reconstructed using `recon-all` [FreeSurfer 6.0.1,RRID:SCR_001847, Dale et al., 1999], and the brain mask estimated previously was refined with a custom variation of the method to reconcile ANTs-derived and FreeSurfer-derived segmentations of the cortical gray-matter of Mindboggle [RRID:SCR_002438, Klein et al., 2017]. Spatial normalization to the ICBM 152 Nonlinear Asymmetrical template version 2009c [Fonov et al., 2009, RRID:SCR_008796] was performed through nonlinear registration with `antsRegistration` [ANTs 2.2.0,RRID:SCR_004757, Avants et al., 2008], using brain-extracted versions of both T1w volume and template. Brain tissue segmentation of cerebrospinal fluid (CSF), white-matter (WM) and gray-matter (GM) was performed on the brain-extracted T1w using `fast` [FSL 5.0.9,RRID:SCR_002823, Zhang et al., 2001].

**Functional data preprocessing** For each of the 4 BOLD runs found per subject (across all tasks and sessions), the following preprocessing was performed. First, a reference volume and its skull-stripped version were generated using a custom methodology of *fMRIPrep*. A deformation field to correct for susceptibility distortions was estimated based on a field map that was co-registered to the BOLD reference, using a custom workflow of *fMRIPrep* derived from D. Greve's `epidewarp.fsl` script and further improvements of HCP Pipelines [Glasser et al., 2013]. Based on the estimated susceptibility distortion, an unwarped BOLD reference was calculated for a more accurate co-registration with the anatomical reference. Head-motion parameters with respect to the BOLD reference (transformation matrices, and six corresponding rotation and translation parameters) are estimated before any spatiotemporal filtering using `mcflirt` [FSL 5.0.9, Jenkinson et al., 2002]. The BOLD time-series (including slice-timing correction when applied) were resampled onto their original, native space by applying a single, composite transform to correct for head-motion and susceptibility distortions. These resampled BOLD time-series will be referred to as *preprocessed BOLD in original space*, or just *preprocessed BOLD*. The BOLD reference was then co-registered to the T1w reference using `bbregister` (FreeSurfer) which implements boundary-based registration [Greve and Fischl, 2009]. Co-registration was configured with nine degrees of freedom to account for distortions remaining in the BOLD reference. The BOLD time-series, were resampled to surfaces on the following spaces: *fsaverage5*. The BOLD time-series were resampled to MNI152NLin2009cAsym standard space, generating a *preprocessed BOLD run in MNI152NLin2009cAsym space*. Several confounding time-series were calculated based on the *preprocessed BOLD*: framewise displacement (FD), DVARS and three region-wise global signals. FD and DVARS are calculated for each functional run, both using their implementations in *Nipype* [following the definitions by Power et al., 2014]. The three global signals are extracted within the CSF, the WM, and the whole-brain masks. Additionally, a set of physiological regressors were extracted to allow for component-based noise correction [*CompCor*, Behzadi et al., 2007]. Principal components are estimated after high-pass filtering the *preprocessed BOLD* time-series (using a discrete cosine filter with 128s cut-off) for the two *CompCor* variants: temporal (tCompCor) and anatomical (aCompCor). Six tCompCor components are then calculated from the top 5% variable voxels within a mask covering the subcortical regions. This subcortical mask is obtained by heavily eroding the brain mask, which ensures it does not include cortical GM regions. For aCompCor, six components are calculated within the intersection of the aforementioned mask and the union of CSF and WM masks calculated in T1w space, after their projection to the native space of each functional run (using the inverse BOLD-to-T1w transformation). The head-motion estimates calculated in the correction step were also placed within the corresponding confounds file. All resamplings can be performed with *a single interpolation step* by composing all the pertinent transformations (i.e. head-motion transform matrices, susceptibility distortion correction when available, and co-registrations to anatomical and template spaces). Gridded (volumetric) resamplings were performed using `antsApplyTransforms` (ANTs), configured with Lanczos interpolation to minimize the smoothing effects of other kernels [Lanczos, 1964]. Non-gridded (surface) resamplings were performed using `mri_vol2surf` (FreeSurfer).

Many internal operations of *fMRIPrep* use *Nilearn* 0.4.2 [Abraham et al., 2014, RRID:SCR_001362], mostly within the functional processing workflow. For more details of the pipeline, see the section corresponding to workflows in *fMRIPrep*'s documentation.

### Original NARPS ex-ante hypotheses

Participating teams will submit yes/no decisions regarding the following anatomical hypotheses for specific contrasts, based on previous results from Tom et al. [2007], De Martino et al. [2010], Canessa et al. [2013], and Canessa et al. [2017].

#### Parametric effect of gain

**1.** Positive effect in ventromedial prefrontal cortex (vmPFC) - for the equal indifference group

**2.** Positive effect in ventromedial PFC - for the equal range group

**3.** Positive effect in ventral striatum - for the equal indifference group

**4.** Positive effect in ventral striatum - for the equal range group

#### Parametric effect of loss

**5.** Negative effect in ventromedial PFC - for the equal indifference group

**6.** Negative effect in ventromedial PFC - for the equal range group

**7.** Positive effect in amygdala - for the equal indifference group

**8.** Positive effect in amygdala - for the equal range group

#### Equal range vs. equal indifference

**9.** Greater positive response to losses in amygdala for equal range condition vs. equal indifference condition.

For each hypothesis, each analysis team would report a binary decision (yes/no) based on a whole-brain correction analysis. Our answers to these hypotheses are found in Table S1.

### Behavioral model

The behavioral model described in the Methods was a logistic regression model which included an intercept and coefficients for losses and gains as predictors (Full model, Supplementary Figure 1). This model was compared to other models to evaluate goodness-of-fit with respect to the Bayesian Information Criterion (BIC). The other models were a model with only an intercept (baseline model, sum total BIC of 33090), a model with an intercept and gains (sum total BIC of 24640), and a model with an intercept and losses (sum total BIC of 26693, see Supplementary Figure 1a). Since the Full model clearly outperformed the other models with overall lower BIC (sum total BIC of 12981; 100% of participants show lower BIC for this model), this model was chosen as the behavioral model that would serve to extract subjective value and decision entropy to calculate the GLMs for the fMRI data.

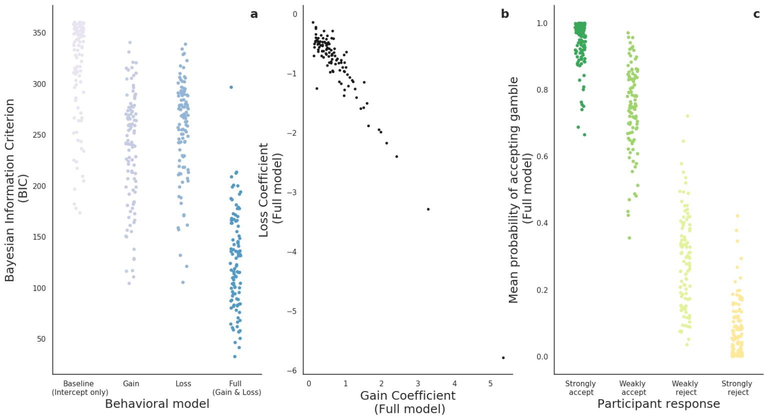


Supplementary Figure 1: Behavioral model selection and evaluation. The Full model is first compared to three other reduced versions (in terms of parameters) with respect to the BIC criterion in **a**. The Full model is evaluated in **b** by plotting the distribution of the losses and gains coefficients. Also, in **c**, the Full model's predictions - mean probability of gamble acceptance - was plotted against the four possible levels of response. Probabilities (model predictions) were averaged across trials for each response level. Not all participants showed responses for all four levels. For all panels, each dot represents a participant.

In Supplementary Figure 1b, we present the distribution of the losses and gains coefficients ($\beta_{losses}$ and $\beta_{gains}$, respectively). There we can observe a very strong correlation between coefficients across participants ($r_{pearson} = -0.96, p < 0.001$). Furthermore, in Supplementary Figure 1c, we evaluate the model predictions of the Full model with respect to the four original (ordered) levels of the dependent variable (participants' response: strongly accept, weakly accept, weakly reject, or strongly reject the gamble). The expected ordering of these levels is reflected in the mean probability of acceptance for the gambles during the experiment: 95.11% (SD = 13.33%), 78.29% (SD = 24.93%), 22.86% (SD = 25.39%) and 7.69% (SD = 16.48%), respectively. All participants presented the expected ordering of the Full model's mean probabilities of gamble acceptance, except for two which showed higher probability for strongly reject than for weakly reject. (Note the expected ordering was adjusted for participants who did not use all four possible responses.)

### Validation of iDE

Due to the relative novelty of inverse decision entropy (iDE) as a proxy variable for confidence, here we validate its use by correlating it with $p_{\text{correct}}$ for the NARPS data (left panel of Supplementary Figure 2), as well as for a two alternative forced choice task from Folke et al. [2017] (middle and right panels of Supplementary Figure 2). For the data from Folke et al. [2017], we also Spearman correlate iDE with explicit confidence ratings (as reported in the main text).

The variable $p_{\text{correct}}$ is another possible proxy variable for confidence previously suggested in De Martino et al. [2013]. It is the subjective probability of being correct, thus closely related to $p_{\text{accept}}$ as defined in the current study:

$$p_{\text{correct}} = \begin{cases} p_{\text{accept}}, & \text{if gamble accepted} \\ p_{\text{reject}}, & \text{otherwise} \end{cases}$$

where $p_{\text{reject}}$ is defined as $1 - p_{\text{accept}}$. It can be similarly defined for a two alternative forced choice task, as in Folke et al. [2017], where one choice option is presented on the left and another one on the right for each trial. For the study in Folke et al. [2017], the items were common retail snacks.

$$p_{\text{correct}} = \begin{cases} p_{\text{choose right item}}, & \text{if } DV \geq 0 \\ p_{\text{choose left item}}, & \text{otherwise} \end{cases}$$

where $p_{\text{choose left item}} = 1 - p_{\text{choose right item}}$ — estimated with a model that outputs probabilities like logistic regression — and $DV$ is the difference in value:

$$DV = v(\text{right item}) - v(\text{left item})$$

The function $v(\cdot)$ assigns a value to each item. Commonly, the values are elicited through a Becker–DeGroot–Marschak (BDM) auction by asking participants to report their willingness-to-pay, usually in units of their monetary currency [Becker et al., 1964]. The variables in the correlation matrix for the middle panel of Supplementary Figure 2 are based on such BDM values — except for the explicit confidence ratings, of course. On the other hand, the right panel of the same figure directly estimates such values (for 32 snack items) by optimizing the log-likelihood of a logistic regression; regressing $DV$ on the choices participants made during the experiment. To our knowledge, directly estimating these values from choices is not common practice and further validates the use of BDM auctions to elicit such values; the correlation between BDM values and the estimated values have a mean Spearman correlation of 0.85 (s.d. $= 0.135$) and were significantly higher than zero ($t(27) = 32.59$, $p < 0.001$). This also provides support for direct estimation of value when BDM values were not collected.

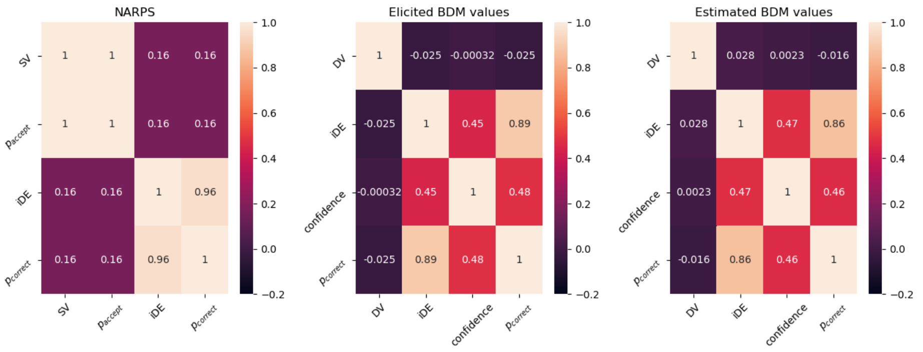


Supplementary Figure 2: Spearman correlation matrices for NARPS and a two alternative forced choice task from Folke et al. [2017]. Left panel shows the (Spearman) correlation matrix for several variables estimated from the NARPS behavioral data: subjective value (SV), $p_{\text{accept}}$, inverse decision entropy (iDE), and $p_{\text{correct}}$, which is the subjective probability of being correct. The middle and right panels are based on behavioral data from Folke et al. [2017] for a similar set of variables: difference in value (DV), iDE, confidence (i.e., explicit ratings), and $p_{\text{correct}}$. The middle panel estimates all variables (except for the confidence ratings) based on explicit reports of willingness-to-pay values for the choice options (i.e., snacks) from a Becker–DeGroot–Marschak (BDM) auction. However, the right panel does not use such BDM values, but estimates them directly from the choices instead. DV is simply the subjective value of an item presented on the right minus the value of an item presented on the left in that experiment.

For completeness, we also present the participant-level plots of iDE against SV in Supplementary Figure 3.

### Loss aversion in the brain

The section on loss aversion serves the function of further validating our behavioral model. It is important to show that a relationship exists between losses and

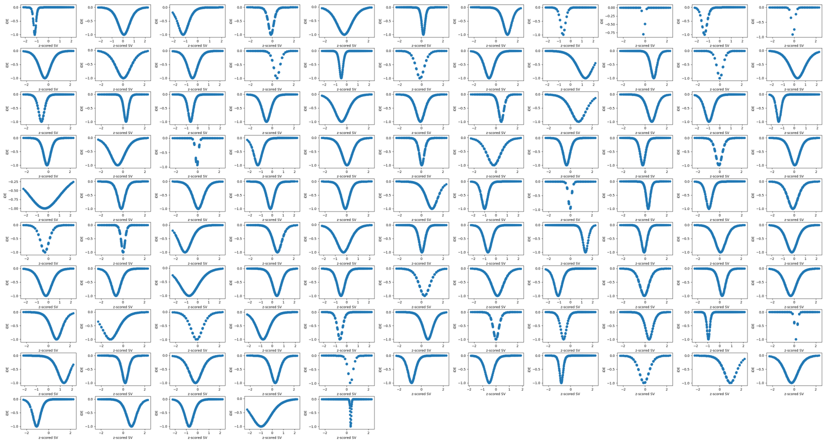

Supplementary Figure 3: Scatterplots of iDE by SV per participant. Each panel represents a participant (104 in total). The y-axis presents inverse decision entropy (iDE) and the x-axis presents (z-scored) subjective value (SV). Each dot represents one choice for a given gamble.

gains in the brain and at the behavioral level. These analyses justify our usage of subjective value as well. As shown in Supplementary Figure 4, behavioral loss aversion correlates with the ratio of voxels that showed sensitivity to losses over sensitivity to gains across participants. Behavioral loss aversion was calculated as the ratio of the loss coefficient divided by the gains coefficient estimated from the behavioral model described in the Methods (M = 1.389, SD = 0.813). (The ratio was multiplied by -1 to keep the relationship to neural activation positive.) To calculate the ratio of loss voxels to gain voxels, we ran a first level GLM on the fMRI data with exactly the same preprocessing and setup as described in the Methods, but instead of using subjective value and decision entropy as parametric modulators we included gains and losses from the experimental design. The first level estimates were averaged with a fixed effects GLM at the second level for each participant. We used a threshold of above or below 2.3 on the $Z$ statistical maps for each participant. Afterwards, we simply counted the number of voxels that survived this thresholding for boths gains and losses for each participant. Finally, the count of loss voxels was divided by the count of gains voxels (M = 1.261, SD = 1.202). To our knowledge, this is a novel way of relating behavioral loss aversion to neural loss aversion.

To assess the relationship between behavioral loss aversion with the losses-to-gains voxel ratio, we ran three different analyses. First, we computed Pearson and Spearman correlations which were both significant (respectively: $r_{pearson} = 0.249, p = 0.011$, and $r_{spearman} = 0.272, p = 0.005$). Second, we ran two robust regressions, downweighting outliers with Huber's loss [Huber et al., 1973], which also showed significant coefficients for: behavioral loss aversion predicting

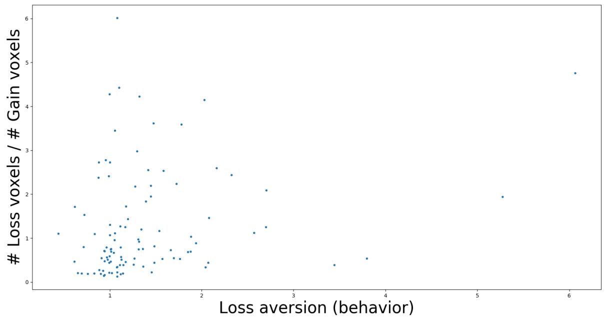


Supplementary Figure 4: Behavioral loss aversion correlates with extent of neural activation. Presents a significant relation between behavioral loss aversion and the ratio of losses to gains voxels across participants.

losses-to-gains voxel ratio ($\beta = 0.373, p = 0.001$) and losses-to-gains voxel ratio predicting behavioral loss aversion ($\beta = 0.07, p = 0.043$). Thirdly, we used a standard outlier detection technique; the standardized difference in fit statistic (DFFITS) [Aguinis et al., 2013] was used with a threshold of $\pm 2\sqrt{(k+1)/n}$, where $k$ is the number of parameters and $n$ is the number of observations, to exclude five outliers. After outlier exclusion, we ran an ordinary least squares (OLS) regression. The same as with the robust regression, both directions of the OLS regression showed significant effects for: behavioral loss aversion ($\beta = 0.516, p = 0.031$) and losses-to-gains voxel ratio ($\beta = 0.092, p = 0.031$).

### Negative main effects of subjective value and inverse decision entropy

Other SV clusters with negative activation include the right occipital fusiform gyrus (33865 voxels, $p < 0.001$, peak $Z = 6.19$), right temporal pole (19881 voxels, $p < 0.001$, peak $Z = 5.53$), left supramarginal gyrus (8058 voxels, $p < 0.001$, peak $Z = 4.75$), right cerebellum (7466 voxels, $p < 0.001$, peak $Z = 5.27$), right frontal pole (7024 voxels, $p < 0.001$, peak $Z = 5.72$), right middle frontal gyrus (4480 voxels, $p < 0.001$, peak $Z = 3.82$), left thalamus (3442 voxels, $p < 0.001$, peak $Z = 4.34$), and vermis crus II (1749 voxels, $p < 0.05$, peak $Z = 4.23$).

For iDE, other clusters include the right and left crus I in the cerebellum (respectively: 29123 voxels, $p < 0.001$, peak $Z = 7.95$, and 27585 voxels, $p < 0.001$, peak $Z = 8.5$), left occipital pole (15664 voxels, $p < 0.001$, peak $Z = 8.28$), right caudate (13001 voxels, $p < 0.001$, peak $Z = 5.19$), right IT (9946 voxels, $p < 0.001$, peak $Z = 7.66$), ventral tegmental area (VTA, 1759 voxels, $p = 0.042$, peak $Z = 4.71$), and right intracalcarine cortex (1713 voxels, $p = 0.049$, peak $Z = 3.78$).

For more information on these clusters see Supplementary Table 1 below.

### Positive main effects of subjective value and inverse decision entropy

Other positively activated clusters for SV include the left precentral and postcentral gyri (2969 voxels, $p = 0.0014$, peak $Z = 6.9$), left inferior temporal (IT) gyrus (2297 voxels, $p = 0.009$, peak $Z = 4.63$), left frontal pole (1983 voxels, $p = 0.023$, peak $Z = 5.62$), left crus I in the cerebellum (1844 voxels, $p = 0.035$, peak $Z = 4.98$), and left lateral occipital cortex (LOC, 1827 voxels, $p = 0.037$, peak $Z = 3.62$).

For iDE, other clusters with positive activation include the left superior frontal gyrus (3852 voxels, $p < 0.001$, peak $Z = 6.38$) and three clusters in the cerebellum: right VIIIb (2255 voxels, $p = 0.009$, peak $Z = 5.08$), left VIIIb (2015 voxels, $p = 0.019$, peak $Z = 4.98$) and left crus II (1752 voxels, $p = 0.043$, peak $Z = 4.87$).

For more information on these clusters see Supplementary Table 1 below.

| | Sign | Cluster Index | Voxels | P | -log10(P) | Z-MAX | Z-MAX X (mm) | Z-MAX Y (mm) | Z-MAX Z (mm) | Z-COG X (mm) | Z-COG Y (mm) | Z-COG Z (mm) | COPE-MAX | COPE-MAX X (mm) | COPE-MAX Y (mm) | COPE-MAX Z (mm) | COPE-MEAN |
|---|---|---|---|---|---|---|---|---|---|---|---|---|---|---|---|---|---|
| Subjective Value | positive | 6 | 17326 | 3.04E-15 | 14.5 | 5.44 | 13 | 15 | -10 | 2.79 | 47.4 | -10.9 | 1.17 | 1 | 63 | -10 | 0.327 |
| | | 5 | 2969 | 0.00138 | 2.86 | 6.9 | -48 | -15 | 52 | -47.7 | -15.8 | 48.8 | 1.56 | -49 | -15 | 58 | 0.632 |
| | | 4 | 2297 | 0.00901 | 2.05 | 4.63 | -54 | -52 | -22 | -53.9 | -51.8 | -18 | 0.663 | -57 | -59 | -24 | 0.319 |
| | | 3 | 1983 | 0.023 | 1.64 | 5.62 | -19 | 44 | 41 | -21.8 | 38.4 | 43.9 | 0.674 | -21 | 45 | 43 | 0.341 |
| | | 2 | 1844 | 0.0352 | 1.45 | 4.98 | -43 | -69 | -41 | -46 | -69.9 | -42.4 | 0.58 | -43 | -76 | -43 | 0.332 |
| | | 1 | 1827 | 0.0371 | 1.43 | 3.62 | -55 | -71 | 34 | -48.6 | -73.8 | 26.4 | 0.959 | -53 | -72 | 32 | 0.41 |
| | negative | 9 | 150923 | 0 | 69 | 8.39 | -44 | -27 | 61 | -28.1 | -25 | 37.5 | 3.04 | -33 | -23 | 70 | 0.453 |
| | | 8 | 33865 | 9.70E-25 | 24 | 6.19 | 31 | -69 | -20 | 32 | -77.7 | -11.8 | 1.36 | 25 | -101 | -13 | 0.387 |
| | | 7 | 19881 | 7.30E-17 | 16.1 | 5.53 | 54 | 13 | -6 | 55.8 | -9.78 | 13.9 | 1.1 | 57 | 19 | -7 | 0.363 |
| | | 6 | 8058 | 1.66E-08 | 7.78 | 4.75 | -33 | 37 | 27 | -30 | 45.2 | 25.2 | 1.04 | -29 | 59 | 25 | 0.357 |
| | | 5 | 7466 | 5.96E-08 | 7.22 | 5.27 | 23 | -60 | -57 | 25.4 | -55.1 | -56.4 | 0.509 | 21 | -59 | -55 | 0.215 |
| | | 4 | 7024 | 1.19E-07 | 6.92 | 5.72 | 33 | 48 | 28 | 29.4 | 51.5 | 19.9 | 0.631 | 33 | 53 | 27 | 0.295 |
| | | 3 | 4480 | 3.15E-05 | 4.5 | 3.82 | 52 | 25 | 30 | 52.6 | 11.5 | 33.4 | 1.14 | 57 | 15 | 35 | 0.38 |
| | | 2 | 3442 | 0.000398 | 3.4 | 4.34 | -13 | -14 | 8 | -13.4 | -19.4 | 6.93 | 0.362 | -11 | -15 | 7 | 0.164 |
| | | 1 | 1749 | 0.0473 | 1.33 | 4.23 | 1 | -77 | -27 | 2.21 | -70.5 | -19.5 | 0.397 | 3 | -77 | -25 | 0.19 |
| Inverse Decision Entropy | positive | 5 | 515033 | 0 | 161 | 8.75 | 6 | 56 | -20 | 1.55 | -28.4 | 10.3 | 44.5 | 20 | -93 | 29 | 8.82 |
| | | 4 | 3852 | 0.000122 | 3.92 | 6.38 | -22 | 28 | 36 | -24.3 | 30.7 | 37.2 | 17.5 | -23 | 31 | 39 | 7.74 |
| | | 3 | 2255 | 0.00918 | 2.04 | 5.08 | 15 | -45 | -56 | 14.5 | -48 | -55.3 | 7.04 | 15 | -45 | -56 | 4 |
| | | 2 | 2015 | 0.0189 | 1.72 | 4.98 | -24 | -42 | -53 | -23.1 | -47.7 | -56.3 | 6.44 | -31 | -45 | -55 | 4 |
| | | 1 | 1752 | 0.0431 | 1.37 | 4.87 | -18 | -77 | -41 | -24.1 | -80.4 | -39.4 | 10.5 | -23 | -79 | -40 | 6.53 |
| | negative | 9 | 300573 | 0 | 112 | 10.2 | 50 | -39 | 53 | 1.34 | -3.54 | 34.7 | 71.3 | 49 | -51 | 56 | 15.6 |
| | | 8 | 29123 | 1.67E-22 | 21.8 | 7.95 | 32 | -60 | -36 | 33.8 | -77.4 | -26.9 | 56.9 | 29 | -99 | -11 | 12.3 |
| | | 7 | 27585 | 1.20E-21 | 20.9 | 8.5 | -36 | -68 | -31 | -28.2 | -69.4 | -39.4 | 22.2 | -41 | -71 | -47 | 8.71 |
| | | 6 | 15664 | 2.46E-14 | 13.6 | 8.28 | -27 | -102 | -11 | -33.7 | -84.5 | -14 | 59 | -29 | -101 | -10 | 16.6 |
| | | 5 | 13001 | 1.80E-12 | 11.7 | 5.19 | 15 | 12 | 11 | 1.62 | -0.729 | 6.37 | 8.02 | 5 | 10 | 5 | 4.16 |
| | | 4 | 9946 | 3.63E-10 | 9.44 | 7.66 | 57 | -41 | -17 | 58.9 | -43.3 | -19.6 | 28.3 | 63 | -51 | -19 | 10.7 |
| | | 3 | 3438 | 0.000348 | 3.46 | 5.86 | -2 | -27 | 26 | 1.45 | -20 | 25 | 15.3 | 1 | -29 | 25 | 5.53 |
| | | 2 | 1759 | 0.0421 | 1.38 | 4.71 | 0 | -16 | -15 | 0.423 | -15.9 | -12.3 | 6.92 | 1 | -21 | -7 | 3.79 |
| | | 1 | 1713 | 0.0488 | 1.31 | 3.78 | 13 | -74 | 11 | 13.3 | -69.8 | 7.53 | 12.4 | 13 | -73 | 10 | 6.52 |

Supplementary Table 1. Maximum *Z* statistics, *p* values and coordinates for cluster activations of all the main effects: positive effect of subjective value, negative effect of subjective value, positive effect of inverse decision entropy, and negative effect of inverse decision entropy. Coordinates are in millimeters (mm) for MNI152. COG: center of gravity and COPE: contrasts of parameter estimates.

### Conjunction of subjective value and inverse decision entropy

Other clusters for conjunction of negative effects include left and right LOC/occipital pole (respectively: 12027 voxels, $p < 0.001$, peak $Z = 4.61$, and 6155 voxels, $p < 0.001$, peak $Z = 5.21$), left precentral gyrus (8099 voxels, $p < 0.001$, peak $Z = 4.51$), right inferior and middle frontal gyrus (respectively, IFG: 4477 voxels, $p < 0.001$, peak $Z = 4.75$ and MFG: 2503 voxels, $p = 0.028$, peak $Z = 3.82$).

For the conjunction of negative SV with positive iDE, there was another observed cluster in the right occipital fusiform gyrus (2417 voxels, $p = 0.035$, peak $Z = 4.41$). We provide extra details on the mentioned clusters: left postcentral gyrus (5216 voxels, $p < 0.001$, peak $Z = 4.54$), right LOC (3359 voxels, $p = 0.003$, peak $Z = 4.84$), and cingulate gyrus (2638 voxels, $p = 0.019$, peak $Z = 4.18$).

For more information on these clusters see Supplementary Table 2 below.

| Subjective Value | Inverse Decision Entropy | Cluster Index | Voxels | P | -log10(P) | Z-MAX | Z-MAX X (mm) | Z-MAX Y (mm) | Z-MAX Z (mm) | Z-COG X (mm) | Z-COG Y (mm) | Z-COG Z (mm) |
|---|---|---|---|---|---|---|---|---|---|---|---|---|
| + | + | 1 | 14732 | 2.70E-12 | 11.6 | 5.02 | 7 | 51 | -20 | 88.4 | 176 | 60.1 |
| negative | negative | 7 | 25820 | 7.53E-19 | 18.1 | 5.76 | -47 | -24 | 61 | 121 | 91.2 | 127 |
| | | 6 | 14195 | 6.09E-12 | 11.2 | 4.93 | -4 | 17 | 37 | 87.6 | 137 | 123 |
| | | 5 | 12027 | 1.82E-10 | 9.74 | 4.61 | 30 | -90 | -19 | 59.8 | 40.4 | 58.6 |
| | | 4 | 8099 | 1.79E-07 | 6.75 | 4.51 | -56 | 1 | 36 | 139 | 142 | 80.2 |
| | | 3 | 6155 | 6.38E-06 | 5.2 | 5.21 | -35 | -90 | -15 | 119 | 32.5 | 57.8 |
| | | 2 | 4477 | 2.23E-04 | 3.65 | 4.75 | 54 | 22 | -3 | 41.7 | 147 | 71.5 |
| | | 1 | 2503 | 0.0277 | 1.56 | 3.82 | 52 | 25 | 30 | 38.2 | 142 | 107 |
| | positive | 6 | 15390 | 1.01E-12 | 12 | 5.06 | -64 | -38 | 25 | 149 | 84.1 | 91.2 |
| | | 5 | 8805 | 5.96E-08 | 7.22 | 4.55 | 68 | -44 | 33 | 27.6 | 96.1 | 93.6 |
| | | 4 | 5216 | 4.46E-05 | 4.35 | 4.54 | -25 | -40 | 71 | 106 | 77.3 | 138 |
| | | 3 | 3359 | 0.00307 | 2.51 | 4.84 | 45 | -81 | 12 | 48.5 | 43.1 | 81.3 |
| | | 2 | 2638 | 0.0193 | 1.71 | 4.18 | -2 | -18 | 44 | 90.3 | 115 | 113 |
| | | 1 | 2417 | 0.0349 | 1.46 | 4.41 | 27 | -74 | -16 | 64.2 | 57.5 | 56.6 |

Supplementary Table 2. Maximum *Z* statistics, *p* values and coordinates for cluster activations of all the conjunctions of subjective value and inverse decision entropy: positive effect (+) of subjective value with positive (+) effect of inverse decision entropy, negative effect of both variables, and negative effect of subjective value with positive effect of decision entropy. Coordinates are in millimeters (mm) for MNI152. COG: center of gravity.

### Contrast of subjective value and inverse decision entropy

We observed cerebellar clusters where inverse decision entropy showed a stronger **negative** effect than subjective value, which include: left and right Crus I (respectively: 4265 voxels, $p < 0.001$, peak $Z = 3.35$, and 4575 voxels, $p < 0.001$, peak $Z = 3.35$), and left Crus II (2869 voxels, $p = 0.01$, peak $Z = 3.15$).

Also, other clusters where subjective value had a significantly larger **negative** effect than inverse decision entropy included: left orbitofrontal cortex (3561 voxels, $p < 0.001$, peak $Z = 3.54$), right frontal pole (3093 voxels, $p < 0.001$, peak $Z = 3.54$), left postcentral gyrus (2719 voxels, $p < 0.001$, peak $Z = 3.54$), right precentral gyrus (2522 voxels, $p = 0.026$, peak $Z = 3.54$), left frontal pole (2470 voxels, $p = 0.030$, peak $Z = 3.54$), and right LOC (2397 voxels, $p = 0.037$, peak $Z = 3.54$).

For more information on these clusters see Supplementary Table 3 below.

| | Sign | Cluster Index | Voxels | P | -log10(P) | Z-MAX | Z-MAX X (mm) | Z-MAX Y (mm) | Z-MAX Z (mm) | Z-COG X (mm) | Z-COG Y (mm) | Z-COG Z (mm) |
|---|---|---|---|---|---|---|---|---|---|---|---|---|
| Subjective Value | negative | 7 | 29900 | 5.23E-21 | 20.3 | 3.54 | -7 | 18 | 30 | -22 | -21 | 59 |
| | | 6 | 3561 | 1.88E-03 | 2.73 | 3.54 | -37 | 21 | -19 | -49 | 23 | -6 |
| | | 5 | 3093 | 5.96E-03 | 2.23 | 3.54 | 21 | 58 | 11 | 29 | 52 | 21 |
| | | 4 | 2719 | 1.56E-02 | 1.81 | 3.54 | -59 | -14 | 28 | -55 | -18 | 21 |
| | | 3 | 2522 | 0.0263 | 1.58 | 3.54 | 28 | -12 | 49 | 30 | -7 | 61 |
| | | 2 | 2470 | 0.0302 | 1.52 | 3.54 | -28 | 46 | 16 | -26 | 53 | 21 |
| | | 1 | 2397 | 0.0368 | 1.43 | 3.54 | 33 | -88 | -21 | 36 | -81 | -18 |
| Inverse Decision Entropy | negative | 7 | 154059 | 0 | 66.2 | 3.54 | -45 | 44 | -23 | -16 | -19 | 35 |
| | | 6 | 106855 | 0 | 51.3 | 3.54 | 44 | 48 | -22 | 27 | 26 | 32 |
| | | 5 | 10039 | 4.91E-09 | 8.31 | 3.54 | 34 | -89 | -21 | 29 | -94 | -8 |
| | | 4 | 7920 | 2.38E-07 | 6.62 | 3.54 | -30 | -94 | -23 | -28 | -96 | -13 |
| | | 3 | 4575 | 0.00018 | 3.75 | 3.35 | 38 | -53 | -39 | 36 | -60 | -34 |
| | | 2 | 4265 | 0.00036 | 3.44 | 3.35 | -25 | -66 | -37 | -34 | -64 | -33 |
| | | 1 | 2869 | 0.0106 | 1.98 | 3.15 | -33 | -78 | -52 | -38 | -70 | -50 |
| | + | 1 | 311318 | 0 | 108 | 3.54 | -44 | -15 | -36 | 0 | -35 | 12 |
| Difference in absolute value | | 5 | 9517 | 1.25E-06 | 5.9 | 5.79 | 21 | -57 | 19 | 9 | -66 | 31 |
| | | 4 | 8699 | 5.36E-06 | 5.27 | 5.4 | 43 | 26 | 38 | 40 | 27 | 39 |
| | | 3 | 5750 | 0.00151 | 2.82 | 5.68 | -61 | -60 | 12 | -54 | -52 | 12 |
| | | 2 | 5236 | 0.00443 | 2.35 | 5.38 | -33 | -57 | 62 | -39 | -47 | 60 |
| | | 1 | 4321 | 0.0325 | 1.49 | 5.42 | 39 | -79 | 38 | 39 | -62 | 50 |

Supplementary Table 3. Maximum *Z* statistics, *p* values and coordinates for cluster activations of all the contrasts of subjective value and inverse decision entropy, both signed and unsigned (absolute value). Negative effect of subjective value, negative effect of inverse decision entropy, positive (+) effect of inverse decision entropy, and difference in absolute value. Coordinates are in millimeters (mm) for MNI152. COG: center of gravity.

### Correlations of voxelwise $Z$ statistics between subjective value and inverse decision entropy

Here, we present the correlations of the $Z$ statistics between SV and iDE. These statistics have the added advantage of incorporating the variance across subjects for each voxel — even though all statistics have been estimated with FSL's mixed effects model (FLAME 1). However, our conclusions remain the same. Frontal medial cortex shows the strongest correlation, $r = 0.616, p < 0.001$, and the correlation remains positive at the whole brain level, $r = 0.295, p < 0.001$. Both left NA, $r = 0.439, p < 0.001$, and right NA, $r = 0.495, p < 0.001$, show strong correlations between SV and iDE as well, followed by the left amygdala, , $r = 0.134, p < 0.001$. With this analysis we only see a change in correlation for the right amygdala; whereas the beta weights showed a Spearman correlation of 0.281 ($p < 0.001$) the correlation of SV and iDE $Z$ statistics is dramatically reduced, $r = 0.030, p = 0.002$.

### Median split of SV

To address concerns that the results may be sensitive to effects of valence within SV (negative domain vs. positive domain), we ran a GLM of the fMRI data in the same way as described in the main text but splitting SV on participant specific medians. We do not observe major differences in the areas that were activated but we do observe that the sign of the betas change depending on whether SV is below the median (middle column of Supplementary Figure 5) or above the median (left column of Supplementary Figure 5). When SV is above the median, it correlates positively with iDE: whole-brain Spearman correlation of 0.173 between betas ($p < 0.001$) and 0.169 between $Z$ statistics ($p < 0.001$). The sign is inverted when SV is below the median: whole-brain Spearman correlation of -0.144 between betas ($p < 0.001$) and -0.142 between $Z$ statistics ($p < 0.001$). This is consistent with our argument that iDE better describes activations in the brain given that iDE is lower when SV is closer to the median. We can conclude that splitting SV in this manner is analogous to explicitly modelling the quadratic relation of SV to iDE that is mentioned in the main text. We also do not observe major changes for activations of iDE (right column of Supplementary Figure 5) when compared to the results reported in the main text.

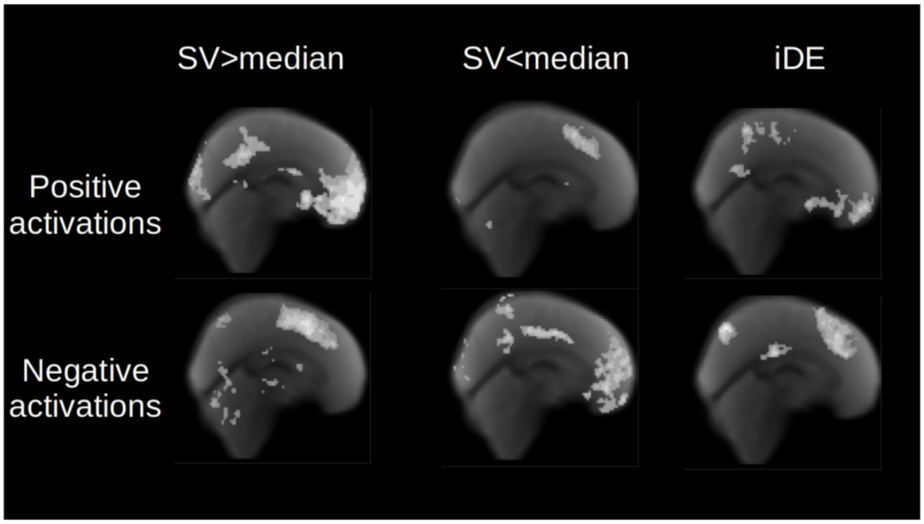


Supplementary Figure 5: Positive and negative activations of median split SV and iDE on the medial surface. The left and middle columns show activations for SV above the participant specific median and below the participant specific median, respectively. The right column shows activations for iDE. The top row presents positive activations and the bottom row presents negative activations.

Joke Durnez, Jasper Degryse, Beatrijs Moerkerke, Ruth Seurinck, Vanessa Sochat, Russell Poldrack, and Thomas Nichols. Power and sample size calculations for fMRI studies based on the prevalence of active peaks. *bioRxiv*, page 49429, 2016.

Oscar Esteban, Ross Blair, Christopher J. Markiewicz, Shoshana L. Berleant, Craig Moodie, Feilong Ma, Ayse Ilkay Isik, Asier Erramuzpe, Mathias Kent, James D. andGoncalves, Elizabeth DuPre, Kevin R. Sitek, Daniel E. P. Gomez, Daniel J. Lurie, Zhifang Ye, Russell A. Poldrack, and Krzysztof Gorgolewski. fmriprep. *Software*, 2018a. doi: 10.5281/zenodo.852659.

Oscar Esteban, Christopher Markiewicz, Ross W Blair, Craig Moodie, Ayse Ilkay Isik, Asier Erramuzpe Aliaga, James Kent, Mathias Goncalves, Elizabeth DuPre, Madeleine Snyder, Hiroyuki Oya, Satrajit Ghosh, Jessey Wright, Joke Durnez, Russell Poldrack, and Krzysztof Gorgolewski. fMRIPrep: a robust preprocessing pipeline for functional MRI. *Nature Methods*, 2018b. doi: 10.1038/s41592-018-0235-4.

Tomas Folke, Catrine Jacobsen, Stephen M Fleming, and Benedetto De Martino. Explicit representation of confidence informs future value-based decisions. *Nature Human Behaviour*, 1(1):2, 2017.

VS Fonov, AC Evans, RC McKinstry, CR Almli, and DL Collins. Unbiased nonlinear average age-appropriate brain templates from birth to adulthood. *NeuroImage*, 47, Supplement 1:S102, 2009. doi: 10.1016/S1053-8119(09)70884-5.

Matthew F. Glasser, Stamatios N. Sotiropoulos, J. Anthony Wilson, Timothy S. Coalson, Bruce Fischl, Jesper L. Andersson, Junqian Xu, Saad Jbabdi, Matthew Webster, Jonathan R. Polimeni, David C. Van Essen, and Mark Jenkinson. The minimal preprocessing pipelines for the human connectome project. *NeuroImage*, 80:105–124, 2013. ISSN 1053-8119. doi: 10.1016/j.neuroimage.2013.04.127. URL http://www.sciencedirect.com/science/article/pii/S1053811913005053.

K. Gorgolewski, C. D. Burns, C. Madison, D. Clark, Y. O. Halchenko, M. L. Waskom, and S. Ghosh. Nipype: a flexible, lightweight and extensible neuroimaging data processing framework in python. *Frontiers in Neuroinformatics*, 5:13, 2011. doi: 10.3389/fninf.2011.00013.

Krzysztof Gorgolewski, Tibor Auer, Vince D Calhoun, R Cameron Craddock, Samir Das, Eugene P Duff, Guillaume Flandin, Satrajit S Ghosh, Tristan Glatard, Yaroslav O Halchenko, and others. The brain imaging data structure, a format for organizing and describing outputs of neuroimaging experiments. *Scientific Data*, 3:160044, 2016.

Krzysztof Gorgolewski, Oscar Esteban, Christopher J. Markiewicz, Erik Ziegler, David Gage Ellis, Michael Philipp Notter, Dorota Jarecka, Hans Johnson, Christopher Burns, Alexandre Manhães-Savio, Carlo Hamalainen, Benjamin Yvernault, Taylor Salo, Kesshi Jordan, Mathias Goncalves, Michael Waskom, Daniel Clark, Jason Wong, Fred Loney, Marc Modat, Blake E Dewey, Cindee Madison, Matteo Visconti di Oleggio Castello, Michael G. Clark, Michael Dayan, Dav Clark, Anisha Keshavan, Basile Pinsard, Alexandre Gramfort, Shoshana Berleant, Dylan M. Nielson, Salma Bougacha, Gael Varoquaux, Ben Cipollini, Ross Markello, Ariel Rokem, Brendan Moloney, Yaroslav O. Halchenko, Demian Wassermann, Michael Hanke, Christian Horea, Jakub Kaczmarzyk, Gilles de Hollander, Elizabeth DuPre, Ashley Gillman, David Mordom, Colin Buchanan, Rosalia Tungaraza, Wolfgang M. Pauli, Shariq Iqbal, Sharad Sikka, Matteo Mancini, Yannick Schwartz, Ian B. Malone, Mathieu Dubois, Caroline Frohlich, David Welch, Jessica Forbes, James Kent, Aimi Watanabe, Chad Cumba, Julia M. Huntenburg, Erik Kastman, B. Nolan Nichols, Arman Eshaghi, Daniel Ginsburg, Alexander Schaefer, Benjamin Acland, Steven Giavasis, Jens Kleesiek, Drew Erickson, René Küttner, Christian Haselgrove, Carlos Correa, Ali Ghayoor, Franz Liem, Jarrod Millman, Daniel Haehn, Jeff Lai, Dale Zhou, Ross Blair, Tristan Glatard, Mandy Renfro, Siqi Liu, Ari E. Kahn, Fernando Pérez-García, William Triplett, Leonie Lampe, Jörg Stadler, Xiang-Zhen Kong, Michael Hallquist, Andrey Chetverikov, John Salvatore, Anne Park, Russell Poldrack, R. Cameron Craddock, Souheil Inati, Oliver Hinds, Gavin Cooper, L. Nathan Perkins, Ana Marina, Aaron Mattfeld, Maxime Noel, Lukas Snoek, K Matsubara, Brian Cheung, Simon Rothmei, Sebastian Urchs, Joke Durnez, Fred Mertz, Daniel Geisler, Andrew Floren, Stephan Gerhard, Paul Sharp, Miguel Molina-Romero, Alejandro Weinstein, William Broderick, Victor Saase, Sami Kristian Andberg, Robbert Harms, Kai Schlamp, Jaime Arias, Dimitri Papadopoulos Orfanos, Claire Tarbert, Arielle Tambini, Alejandro De La Vega, Thomas Nickson, Matthew Brett, Marcel Falkiewicz, Kornelius Podranski, Janosch Linkersdörfer, Guillaume Flandin, Eduard Ort, Dmitry Shachnev, Daniel McNamee, Andrew Davison, Jan Varada, Isaac Schwabacher, John Pellman, Martin Perez-Guevara, Ranjeet Khanuja, Nicolas Pannetier, Conor McDermottroe, and Satrajit Ghosh. Nipype. *Software*, 2018. doi: 10.5281/zenodo.596855.

Douglas N Greve and Bruce Fischl. Accurate and robust brain image alignment using boundary-based registration. *NeuroImage*, 48(1):63–72, 2009. ISSN 1095-9572. doi: 10.1016/j.neuroimage.2009.06.060.

Peter J Huber et al. Robust regression: asymptotics, conjectures and monte carlo. *The Annals of Statistics*, 1(5):799–821, 1973.

Mark Jenkinson, Peter Bannister, Michael Brady, and Stephen Smith. Improved optimization for the robust and accurate linear registration and motion correction of brain images. *NeuroImage*, 17(2):825–841, 2002. ISSN 1053-8119. doi: 10.1006/nimg.2002.1132. URL http://www.sciencedirect.com/science/article/pii/S1053811902911328.

Arno Klein, Satrajit S. Ghosh, Forrest S. Bao, Joachim Giard, Yrjö Häme, Eliezer Stavsky, Noah Lee, Brian Rossa, Martin Reuter, Elias Chaibub Neto, and Anisha Keshavan. Mindboggling morphometry of human brains. *PLOS Computational Biology*, 13(2):e1005350, 2017. ISSN 1553-7358. doi: 10.1371/journal.pcbi.1005350. URL http://journals.plos.org/ploscompbiol/article?id=10.1371/journal.pcbi.1005350.

C. Lanczos. Evaluation of noisy data. *Journal of the Society for Industrial and Applied Mathematics Series B Numerical Analysis*, 1(1):76–85, 1964. ISSN 0887-459X. doi: 10.1137/0701007. URL http://epubs.siam.org/doi/10.1137/0701007.

Xiangrui Li, Paul S Morgan, John Ashburner, Jolinda Smith, and Christopher Rorden. The first step for neuroimaging data analysis: DICOM to NIfTI conversion. *Journal of neuroscience methods*, 264:47–56, 2016.

Denis G Pelli. The VideoToolbox software for visual psychophysics: Transforming numbers into movies. *Spatial vision*, 10(4):437–442, 1997.

Jonathan D. Power, Anish Mitra, Timothy O. Laumann, Abraham Z. Snyder, Bradley L. Schlaggar, and Steven E. Petersen. Methods to detect, characterize, and remove motion artifact in resting state fmri. *NeuroImage*, 84(Supplement C):320–341, 2014. ISSN 1053-8119. doi: 10.1016/j.neuroimage.2013.08.048. URL http://www.sciencedirect.com/science/article/pii/S1053811913009117.

Sabrina M Tom, Craig R Fox, Christopher Trepel, and Russell A Poldrack. The neural basis of loss aversion in decision-making under risk. *Science*, 315(5811): 515–518, 2007.

N. J. Tustison, B. B. Avants, P. A. Cook, Y. Zheng, A. Egan, P. A. Yushkevich, and J. C. Gee. N4itk: Improved n3 bias correction. *IEEE Transactions on Medical Imaging*, 29(6):1310–1320, 2010. ISSN 0278-0062. doi: 10.1109/TMI.2010.2046908.

Y. Zhang, M. Brady, and S. Smith. Segmentation of brain MR images through a hidden markov random field model and the expectation-maximization algorithm. *IEEE Transactions on Medical Imaging*, 20(1):45–57, 2001. ISSN 0278-0062. doi: 10.1109/42.906424.
